# Functional characterization of ovine dorsal root ganglion neurons reveals peripheral sensitization after osteochondral defect

**DOI:** 10.1101/2021.02.26.432434

**Authors:** Sampurna Chakrabarti, Minji Ai, Wai Ying Katherine Wong, Karin Newell, Frances M.D. Henson, Ewan St. John Smith

## Abstract

**Objective:** Knee joint trauma can cause an osteochondral defect (OD), a risk factor for osteoarthritis and cause of debilitating pain in patients. Modelling OD in rodents is difficult due to their smaller joint size. This study proposes sheep as a translationally relevant model to understand the neuronal basis of OD pain.

**Methods:** Unilateral 6 mm deep OD was induced in adult sheep, 2-6 weeks after which dorsal root ganglion neurons (DRG neurons) were cultured from the control and OD side. Functional assessment of neuronal excitability and activity of the pain-related ion channels, TRPV1 and P2X3, was carried out using electrophysiology and Ca^2+^-imaging. Immunohistochemistry was utilized to verify expression of pain-related proteins.

**Results:** An increased proportion of OD DRG neurons (sheep, n = 3, Ctrl neurons, n =15, OD neurons, n = 16) showed spontaneous electrical excitability (p = 0.009, unpaired t-test) and hyperexcitability upon TRPV1 agonist (capsaicin) application (p = 0.04, chi-sq test). Capsaicin also produced Ca^2+^ influx in an increased proportion of OD DRG neurons isolated (p = 0.001, chi-sq test). By contrast, neither protein expression, nor functionality of the P2X3 ion channel were altered in OD neurons.

**Conclusions:** We provide evidence of increased excitability of DRG neurons (which is an important neural correlate of pain) and TRPV1 function in an OD sheep model. Our data show that functional assessment of sheep DRG neurons can provide important insights into the neural basis of OD pain and thus potentially prevent its progression into arthritic pain.

## Introduction

Osteochondral lesions are detected in ∼60% of patients who undergo knee arthroscopies^1^. Clinically, osteochondral defects (OD) constitute damage to bones (osteo) and cartilage (chondral) and commonly present as pain and swelling of joints after an acute injury, initial radiographs often being negative for lesions^2^. OD is diagnosed only if pain on weight bearing persists for more than 4-6 weeks post-injury and can also reduce the quality of life of patients to a similar extent to individuals with late stage osteoarthritis (OA)^1,2^. Indeed, OD in adults is a risk factor for progression to OA, highlighting the importance of OD research as prevention for OA progression^3^. In OD, pain is suspected to arise from hyperexcitability of sensory nerves innervating the subchondral bone which is further amplified by secretion of inflammatory mediators from the aneural articular cartilage and synovial membrane^3^. However, direct evidence of sensory nerve hyperexcitability in OD and the mechanisms involved in such peripheral sensitization is lacking.

Peripheral sensitization is commonly studied by electrophysiological recordings of isolated dorsal root ganglion neurons (DRG, location of the cell bodies of sensory nerves innervating the body, but not the head) harvested from rodents. However, rodent models of OD are difficult to create and less translatable because of their smaller cartilage volume (rats: 2.17 mm^3^) compared to humans (552 mm^3^)^4^. The thinner cartilage in rodents restricts the amount of damage that can be induced, and therefore is less translatable to the human disease. In contrast, large animals, such as sheep, have a similar cartilage volume (359 mm^3^) to humans and OD can successfully be induced in their joints as evidenced by histological scoring and reduced activity^4,5^. Consequently, the majority of OD research is focused on developing strategies for bone and cartilage regeneration, such as implantation of biomaterials in large animal models^4^. Whether such regeneration strategies also decrease any neuronal sensitization that occurs in OD, and hence pain, is largely unknown due to the lack of expertise in isolation and recording of DRG neurons from large animals.

In this study, we provide the first evidence of peripheral sensitization in an OD model of the sheep stifle joint by electrophysiological recording and Ca^2+^ imaging of isolated DRG neurons. Such an in vitro experimental paradigm could be utilized to identify translatable pain targets for OD and OA and as an outcome measure of future pain therapeutics and cartilage regeneration technologies.

## Material and methods

This study was approved by the University of Cambridge Animal Welfare and Ethical Review Body and the UK Home Office (Project License 70/8165). See supplementary file for detailed methodology.

### Animals

Six skeletally mature female Welsh Mountain sheep (3.2 ±0.8 years, 40-44kg) housed in flocks under natural conditions with the same feed, husbandry and location were included in the study. A 6-mm deep, 8-mm wide OD was created using a hand drill on the left stifle joint of these sheep under anesthesia (thiopentone, 3 mg/kg intravenous, followed by isoflurane), accompanied by analgesia (carprofen, 4 mg/kg, intramuscular) and prophylactic antibiotic administration (procain penicillin). Animals were sacrificed by injection of 40 ml 20% (w/v, intravenous.) pentobarbitone sodium at 2, 4, and 6 weeks post-surgery. A semiquantitative histological analysis using International Cartilage Repair Society macroscopic score was carried out blindly as previously established^5^.

### DRG neuron isolation and culture

L3 – L4 DRG from control and OD sides of sheep were collected after transection of vertebral bodies from the lumbar region in ice cold dissociation media as described before^6,7^. The collected DRG were cut into ∼3 mm^3^ pieces before enzymatic digestion with collagenase followed by trypsin (1mg/ml each). The DRG were then kept in trypsin and collagenase digestion solutions for ∼30 min each, including a 5-10 min shaking step at the beginning of incubation; before mechanical trituration and plating on Poly-D-lysine and laminin coated glass bottomed dishes (MatTek, P35GC-1.5-14-C). Plated neurons were incubated at 37 °C, 5% CO_2_.

### Whole-cell patch clamp electrophysiology

DRG neuron recordings were made following overnight incubation. Tested neurons were bathed in extracellular solution (ECS) and recorded from using an EPC-10 amplifier (HEKA) and Patchmaster© software (HEKA). Glass pipettes with a resistance of 3–6 MΩ loaded with intracellular solution (ICS) were used for whole-cell patch clamp. Action potentials (AP) were recorded under current-clamp mode either without current injection (to record spontaneous firing) or following stepwise current injection (to evoke firing). Transient receptor potential vanilloid receptor 1 (TRPV1) and purinergic (P2X3) ion channel agonists (capsaicin (1 μM) and α,β, mATP (30 μM), Sigma) were applied to DRG neurons for 10s to determine their ability to evoke APs under current clamp mode. Current-voltage relationships were obtained using a standard voltage-step protocol under voltage-clamp mode (Figure 1G).

**Figure 1:**
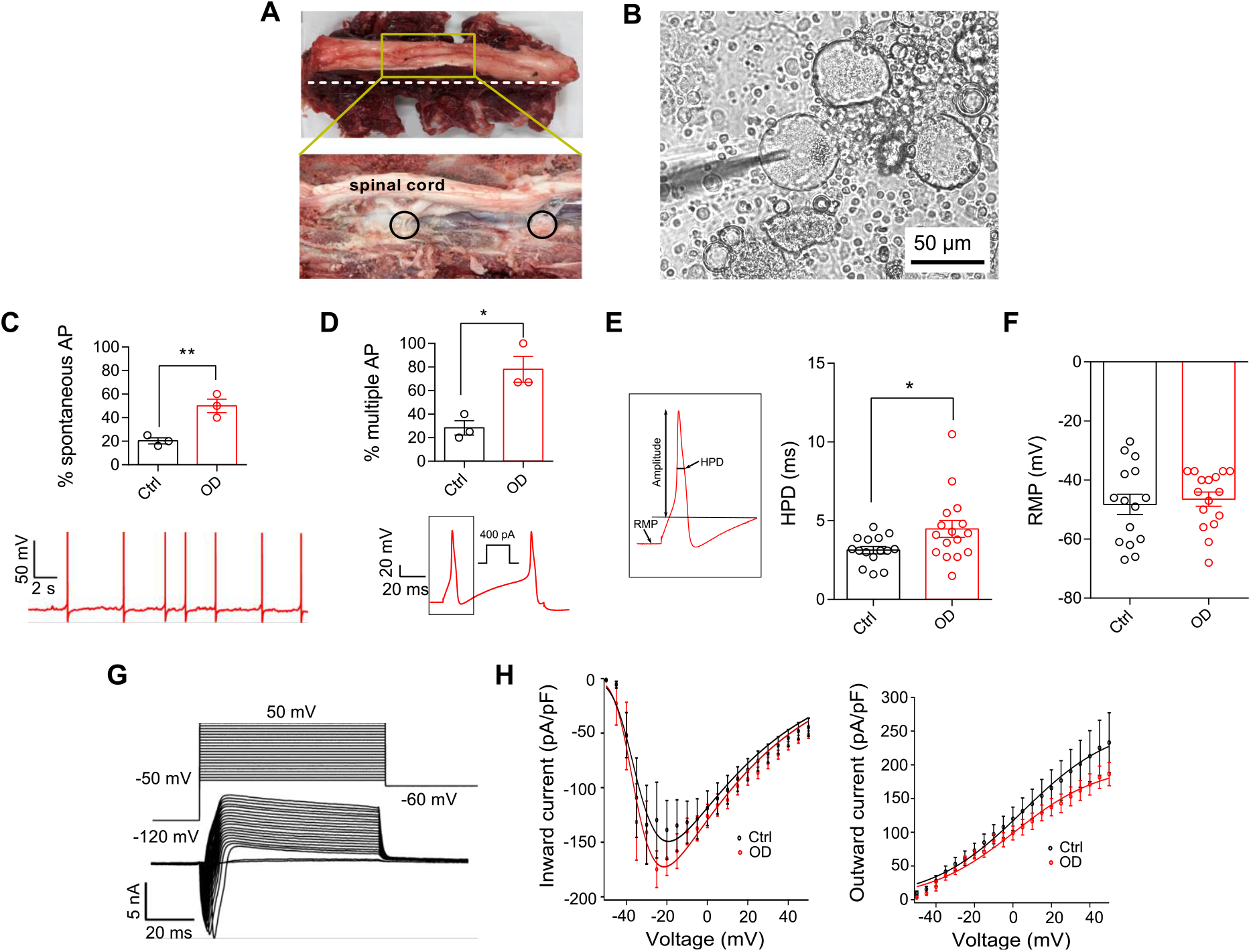
A) Photo showing intact (top) lumbar region of a sheep spine and after transverse section (white dotted dissection line, bottom) to expose DRG (black circles). B) Acutely dissociated sheep DRG neurons in culture. C) Percentage of neurons showing spontaneous activity in ctrl and OD condition (top) and a representative OD neuron with spontaneous activity (bottom). D) Percentage of neurons firing multiple action potentials (AP) upon current injection (top) and a representative trace of multiple firing in response to 400 pA injected current (bottom). E) Schematic representation of AP properties (left) and distribution of half peak duration (HPD) (right) and (F) resting membrane potential (RMP) in ctrl and defect conditions. G) Representative traces of currents evoked from a neuron at different voltages and (H) plots of the inward and outwards currents in ctrl (black) and OD (red) neurons. *: p < 0.05, **: p < 0.01, unpaired t-test.

### Ca^2+^ imaging

DRG neurons were incubated in 10 μM of the Ca^2+^ indicator Fluo-4 AM (Invitrogen, UK) for 30 min at room temperature (21°C). Neurons were then washed with ECS and placed under a microscope (Nikon Eclipse Tie–S, Nikon) for imaging. Fluro-4 fluorescence was excited by a 470 nm LED (Cairn Research) and images were captured by a digital camera (Zyla cSMOS, Andor) at 1 Hz with 50 ms exposure time using Micro-Manager software (v1.4; NIH). 50 mM KCl was used as positive control. Data analysis was conducted using a custom made R toolbox (https://github.com/amapruns/Calcium-Imaging-Analysis-with-R).

### Immunohistochemistry

L3 – L4 DRG from OD and control sides of sheep were collected in Zamboni’s fixative, embedded in Shandon M-1 embedding matrix (Thermo Fisher Scientific, UK), snap frozen and stored at −80°C as described before. One to three 12 μm sections were chosen randomly from both sides for staining as previously established^6^ using an anti-CGRP antibody (1:5000, Sigma C8189, anti-rabbit polyclonal), an anti-P2X3 antibody (1:1000, Alomone APR016, anti-rabbit polyclonal) and an Alexa-488 conjugated secondary antibody (1:1000, Invitrogen A21206). The mean gray value of each DRG neuron was measured in ImageJ and a custom-made R toolkit (https://github.com/amapruns/Immunohistochemistry_Analysis) was used to identify positive neurons with manual validation as previously described^6^.

### Statistics

Data are shown as mean ± Standard Errors of Mean (SEM). Two group comparisons were made with Student’s unpaired t-test, and percentage comparison was done using a chi-square test. One-way ANOVA (Analysis of Variance) with Tukey’s post-hoc test was used to compare the significance among three groups.

## Results

### OD induces hyperexcitability in sheep DRG neurons

A 6 mm OD was created unilaterally on the femoral condyle of sheep (n=3) stifle joint which resulted in cartilage damage (assessed macroscopically, Supplementary Table 1). Pain is the major symptom of OD in humans and we have previously reported that synovial fluid from patients with painful OA can increase the excitability of mouse DRG neurons^8^. Consequently, we hypothesized that the isolated DRG neurons from the OD side of sheep would show hyperexcitability. To test this hypothesis, we harvested, cultured and performed patch-clamp recording on the DRG neurons isolated from the ctrl (n = 15) and OD (n = 16) sides of the sheep (Figure 1A). We recorded from small-medium sized putative nociceptors (<1000 μm^2^ in area; Figure 1B and Supplementary figure 1) and showed that neurons from the OD side have enhanced spontaneous AP firing in the absence of current injection (p = 0.009, unpaired t-test, Figure 1C) and multiple AP firing (p = 0.01, unpaired t-test, Figure 1D) after current injection. These results suggest that OD enhances excitability of nociceptors. Notably, the AP threshold in both ctrl and OD neurons were ∼100 pA (Supplementary Table 2), which is lower than that reported for murine neurons^6^. Additionally, AP half-peak duration (HPD) was increased in neurons isolated from the OD side (p = 0.03, unpaired t-test, Figure 1E) although no change in resting membrane potential (RMP) or other AP properties was observed (Figure 1F, Supplementary Table 2). Increased HPD is suggestive of increased voltage-gated Ca^2+^ (Ca_v_) and Na_v_1.8 channel function^9^, but no difference in inward (mediated by Na_v_ and Ca_v_) or outward (mediated by K_v_) current-voltage relationship was found between ctrl and OD conditions (Figure 1G,H), although recordings of isolated Na_V_, Ca_V_ and K_V_ currents were not conducted. Taken together, our data suggest that 2-6 weeks after OD, sheep DRG neurons show increased excitability, which is a correlate of pain, and this increase in excitability is not due to significant changes in the activity of voltage-gated ion channels.

### Sheep DRG neurons show increased TRPV1 function after OD

In addition to voltage-gated ion channels, increased functionality of algogen-sensing ion channels can cause nociceptor hyperexcitability^10^. We tested agonists of TRPV1 and P2X3 ion channels (capsaicin and α,β me-ATP respectively) and observed that sheep DRG neurons respond to these known algogens by firing AP (Figure 2A) similar to neurons isolated from mouse and human DRG^10^. Additionally, we observed that the proportion of OD neurons firing above baseline upon application of capsaicin was significantly higher than the ctrl neurons, implicating an increase in TRPV1 function (p = 0.04, chi-sq test, Figure 2B). However, α,β me-ATP produced above baseline firing in a similar proportion of neurons isolated from both control and OD sides, thus arguing against a significant role for P2X3 channel in OD pain (Figure 2B). Next we performed Ca^2+^-imaging on these neurons to reveal that after OD, an increased proportion of neurons respond to capsaicin (ctrl, 35/141, OD, 48/109, p = 0.001, chi-sq test, Figure 2Ci,ii), while the number of neurons responding to α,β me-ATP was not significantly different between ctrl and OD groups (ctrl, 24/115, OD, 23/69, Figure 2Di,ii). Congruently, we showed using IHC on sections of whole DRG that the proportion of P2X3+ neurons was similar (∼30 %) in ctrl and OD conditions (Figure 2Ei,ii), however, protein level expression of TRPV1 could not be validated due to the unavailability of a specific antibody for sheep TRPV1 (tested antibodies listed in Supplementary Table 3). Finally, we probed the expression of the pro-nociceptive neuropeptide, CGRP, in DRG sections because an increase in TRPV1 expression can in turn induce production of CGRP^11^. However, the CGRP+ neurons were ∼30% in both OD and ctrl conditions (as observed before^7^) suggesting that the proportion of peptidergic CGRP neurons is unchanged in OD pain (Figure 2Fi,ii).

**Figure 2:**
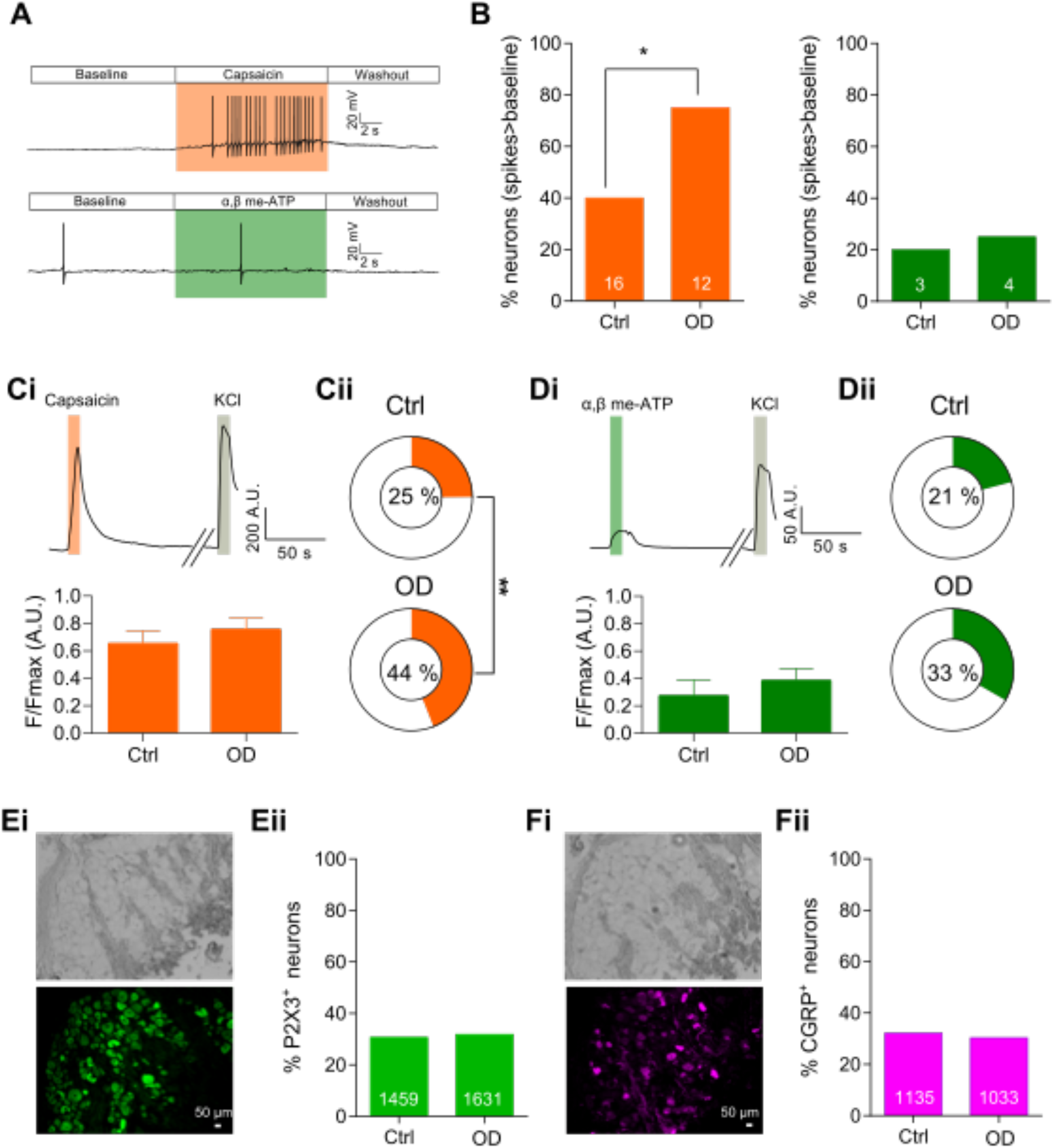
A) Patch-clamp traces showing a representative DRG neuron from the ctrl side firing AP spikes above baseline and same as baseline in response to capsaicin and α,β me-ATP respectively. B) Bar graph showing percentage of neurons with above baseline firing activity in response to capsaicin (left, orange) and α,β me-ATP (right, green). Numbers on the bars represent number of neurons in each condition firing above baseline. Ci) (top) Representative Ca^2+^ trace from a neuron responding to capsaicin and KCl (positive control). (bottom) Magnitude of Ca^2+^ influx in response to capsaicin (ctrl, n = 35, defect, n = 48). Cii) Percentage of neurons responding to capsaicin in each condition. Di) (top) Representative Ca^2+^ trace from a neuron responding to α,β me-ATP and KCl (positive control), (bottom) magnitude of Ca^2+^ influx in response to α,β me-ATP (ctrl, n = 24, defect, n = 23). Dii) Percentage of neurons responding to α,β me-ATP in each condition. Ei) Representative brightfield (top) and anti-P2X3-antibody stained (bottom) image of a whole sheep DRG in cross-section along with the percentage of neurons positive for P2X3 (Eii). Fi) Representative brightfield (top) and anti-CGRP-antibody stained (bottom) image of a whole sheep DRG in cross-section along with the percentage of neurons positive for CGRP (Fii). Numbers on the bars represent neurons positive for the respective antibody staining. *: p < 0.05, **: p < 0.01, chi-sq test.

Taken together our data provide support that OD causes hyperexcitability of sheep DRG neurons and that increased function of TRPV1 is part of the sensitization process. Since TRPV1 is also an important pain target in OA, TRPV1 antagonists utilized for pain-control in OD might also help prevent pain in cases of OD that develop into OA.

## Discussion

We have previously proposed that large animals, such as sheep, can be leveraged as translational models to investigate mechanisms of joint pain in vitro due to their larger joint anatomy and DRG neuron diameter compared to rodents^12^. In this study we provide proof using the ovine OD model (which is difficult to induce in rodents) that it is possible to study neuronal constructs of peripheral sensitization in large animals using tools developed for rodent DRG neurons. For example, we show using whole-cell patch clamp electrophysiology and Ca^2+^-imaging that sheep DRG neurons can be activated by agonists of nociceptive ion channels TRPV1 and P2X3, thus indicating their functional presence.

Importantly, we found using whole-cell patch clamp that an increased proportion of OD neurons fired spontaneous AP, as well as multiple APs within 80 ms in response to injected current. The increase in AP firing was not associated with significant changes in voltage-gated ion channel activity, however, further studies are needed to determine changes in the function of hyperpolarization-activated cyclic nucleotide-gated channels, which have been implicated in sensory nerve firing and pain^13^. We also observed increased AP firing upon administration of the TRPV1 agonist capsaicin in OD neurons, along with an increased proportion of OD neurons responding to capsaicin using Ca^2+^-imaging assay. These data suggest that TRPV1-mediated depolarization can increase AP firing in sensory neurons to a greater extent after OD, which is consistent with mechanistic studies showing a TRPV1-anoctamin 1 interaction that increases prolonged glutamate release to induce pain-related behaviors^14^. Furthermore, our data are consistent with peripheral sensitization observed in models of joint pain^6^, implying that similar nociceptive mechanisms are also at play before induction of arthritis. Therefore, TRPV1 antagonists have the potential to ameliorate or perhaps prevent onset of arthritic pain.

Lastly, we observed that expression of CGRP and P2X3, markers of peptidergic and non-peptidergic sensory neurons respectively, were mostly expressed in small-size DRG neurons (<1000 μm^2^, Supplementary Figure 1) in a similar manner to in rodent and human DRG^15^. Additionally, the lack of change in the percentage of neurons expressing P2X3 correlates with our observation that P2X3 function remained unchanged after OD.

This study highlights that sheep have huge, hitherto untapped, potential in mechanistic joint pain research. However, efforts need to be made to develop tools effective for large animals. For example, due to the unavailability of specific antibodies that work on sheep tissue, we were unable to assess using IHC if enhanced TRPV1 expression occurred in DRG neurons, as well as being unable to investigate if P2X3 and CGRP are coexpressed a subset of DRG neurons as has been observed in human DRG^15^. Nevertheless, our study paves the way for future investigation of pain mechanisms in OD which might help prevent progression of joint trauma to arthritis.

## Contributions

S.C. designed and conducted experiments, analyzed data and wrote the manuscript. M.A. conducted experiments, analyzed data and wrote the paper. K.W. conducted experiments and analyzed data. K.N. conducted experiments and analyzed data. F.M.D.H. and E.S.J.S designed experiments and revised the manuscript. All authors approve the final form of the article.

## Extended Material and methods

This study is approved by the University of Cambridge Animal Welfare and Ethical Review Body (AWERB) and the UK Home Office (Project Licence 70/8165).

### Animals

Six skeletally mature female Welsh Mountain sheep (3.2 ±0.8 years, 40-44kg) were included in the study. Experiments were conducted using sterile surgical procedure carried out by the same surgical team. All animals were housed in flocks outside under natural conditions with the same feed, husbandry and location.

### Animal anesthesia, preparation and surgical technique

Sheep were anaesthetized by intravenous injection of thiopentone (3 mg/kg) and maintained using inhalable anesthetic (a mixture of isoflurane, nitrous oxide, and oxygen). Perioperative analgesia was provided by intramuscular injection of carprofen (4 mg/kg). Antibiotic prophylaxis (procaine penicillin) was given through intramuscular injection. All animals used went through identical surgical procedure under strict aseptic conditions. Each stifle joint was physically examined for abnormality under anesthesia and animals with gross joint instability or pathology was excluded from the study.

The osteochondral defect (OD) was created on the left stifle joint of experimental sheep. Each animal was placed in a dorsal recumbency position following surgical preparation and the left stifle joint was opened through a parapatellar approach. The patellar fat pad was then elevated to access the medial femoral condyle (MFC) where a 6-mm deep, 8-mm wide OD was created using a hand drill. Following surgery, operated animals were kept in small pens for 48h to reduce ambulation prior to allowing them to fully bear weight. Sheep were then housed in large pens or outdoor fields with normal ambulation before being sacrificed by intravenous injection of 40 ml 20% (w/v) pentobarbitone sodium at 2, 4, and 6-weeks post-surgery. Macroscopic scoring of the joints were carried out by F.H. according to the International Cartilage Repair Society guidelines^1^.

### DRG neuron isolation and culture

Dorsal root ganglia (DRG) were dissected from operated sheep immediately after them being sacrificed with slight modifications from previously performed procedures on sheep and other species^2–4^. Sheep were placed on the operating table in the posterior positon and their midline fur was shaved using a veterinary clipper. A surgical scalpel was then used to cut open the skin, retract the obliques and latissimus muscles, after which a bone saw was used to remove the lumbar (L2-L5) part of the vertebrae en-bloc. A dorsal laminectomy was then performed using bone saw to expose the spinal cord and DRG (Figure 1B). Exposed DRG were carefully lifted with a forceps while dissecting scissors were used to simultaneously cut the spinal root and peripheral nerve rami to free the DRG.

L3 – L4 DRG from contralateral and ipsilateral sides of OD sheep were collected and placed in ice cold dissociation media containing L-15 Medium (1X) + GlutaMAX-l (Life Technologies) with 24 mM NaHCO_3_ supplement. Collected DRGs were then transported back to the laboratory on ice for neuronal isolation. Individual DRG were fully immersed in a 35 mm petri dish containing ice cold dissociation media and connective tissues surrounding the DRG were carefully removed using a pair of microscissors. Each DRG was then finely minced into small pieces (3 mm3), submerged in 3 ml collagenase solution (1 mg/ml type I collagenase A (Sigma, UK) with 6 mg/ml Bovine serum albumin (BSA) (Sigma, UK) and placed on a shaker (30 rev/min) for 10 min prior to 15 min incubation in a 37°C incubator. Collagenase solution was then replaced with 3 ml prewarmed trypsin solution (1 mg/ml trypsin (Sigma, UK) with 6 mg/ml BSA in dissociation media) and placed on the shaker for 5 min before 30 min incubation at 37 °C. Enzyme solution was then removed and prewarmed culture media (dissociation medium with 10% (v/v) fetal bovine serum, 2% penicillin/streptomycin and 38 mM glucose) was added to suspend the DRG. Suspended DRG solution was transferred into a 15 ml falcon tube for gentle mechanical trituration with a 1 ml Gilson pipette followed by a brief centrifugation (160 g, 30 s; Biofuge primo, Heraeus Instruments; Hanau, Germany). Supernatant containing dissociated neurons was collected in a fresh tube. This dissociation step was repeated for five times until 10 ml of supernatant was collected. Collected supernatant was centrifuged at 160 g for 5 mins for neuron pelleting, which was then resuspended in 250 μl DRG culture media and plated on Poly-D-lysine and laminin coated glass bottomed dishes (MatTek, P35GC-1.5-14-C) for 3 hours to allow neuron to attach. Additional 2 ml culture media was added to each culture dish following neuron attachment. All neurons were incubated in incubator (37 °C, 5% CO_2_) overnight (8 –10 hours) before electrophysiology and calcium imaging recording.

### Immunohistochemistry

L3 – L4 DRG from contralateral and ipsilateral sides of OD animals were collected as described above. Collected DRG were immediately fixed in Zamboni’s fixative (4% paraformaldehyde and picric acid) for 1 hour and transferred to 30% (w/v) sucrose for overnight incubation at 4°C. Processed DRG were then embedded in Shandon M-1 embedding matrix (Thermo Fisher Scientific, UK), snap frozen in liquid nitrogen and stored at −80°C. Embedded DRG were sectioned by a Leica Cryostat (CM3000; Nussloch, Germany), mounted on Superfrost Plus microscope slides (Thermo Fisher Scientific) and stored at −20 °C until staining. One to three sections were chosen randomly from both operated and non-operated sides for analysis. Staining was carried by following previous established protocol ^5^ and performed blindly by KW. Anti-CGRP antibody (1:5000, Sigma C8189, anti-rabbit polyclonal), and anti-P2X3 antibody (1:1000, Alomone APR016, anti-rabbit polyclonal) following Alexa-488 conjugated secondary antibody (1:1000, Invitrogen A21206, anti-rabbit) and Alexa-568 conjugated secondary antibody (1:500, Invitrogen A-11031, anti-mouse) were used in staining. Mean gray value of each DRG neuron were measured by ImageJ and a custom-made R toolkit (https://github.com/amapruns/Immunohistochemistry_Analysis) was used to score positive neurons with manual validation as previously described ^5^. In brief, a normalized distribution of neurons with the least mean gray value from each section was computed (distribution of minima). All neurons that had a mean gray value greater than 2 standard deviation from the average of the distribution of minima were scored positive.

### Whole cell patch-clamp Electrophysiology

DRG neuron recordings were made following overnight incubation. At least five neurons from OD or non-operated sides of OD animals (n = 3) were recorded. Neurons were bathed in extracellular solution (in mM): NaCl (140), KCl (4), CaCl2 (2), MgCl2 (1), glucose (4), HEPES (10), adjusted to pH 7.4 with NaOH and recorded by EPC-10 amplifier (HEKA) together with Patchmaster© software (HEKA). Glass pipettes used for patching were pulled (P-97, Sutter Instruments) from borosilicate glass capillaries with a resistance of 3–6 MΩ. A ground electrode was placed in the neuron bath to form a closed electric circuit with patching pipette loaded with intracellular solution (in mM): KCl (110), NaCl (10), MgCl_2_ (1), EGTA (1), and HEPES (10), adjusted to pH 7.3 with KOH. Neurons were held at −60 mV with pipette and membrane resistance compensated. Resting membrane potential (RMP), cell resistance and capacitance were recorded on current-clamp mode prior than any testing protocols. Neurons with current evoked action potential and a RMP lower than −40 mV were included for analysis. Neuron images were captured by a 40×objective on microscope (Nikon Eclipse Tie– S, Andor) and a Zyla 5.5 sCMOS camera (Belfast, United Kingdom). Cell area was calculated by ImageJ software following pixel to μm conversion.

Action potentials (AP) were recorded under current-clamp mode without current injection (to record spontaneous firing) following stepwise current injection. Current from 100 pA to 1000 pA was injected for 80 ms through 50 steps and the first evoked AP was analyzed. AP threshold, half peak duration (HPD, ms), amplitude, afterhyperpolarization duration (AHP, ms), and afterhyperpolarization amplitude (AHP, mV), were measured using FitMaster (HEKA) software or IgorPro software (Wavemetrics) as previously described^5^.

Ion channel agonists (capsaicin (1 μM, TRPV1) and α,β, mATP (30 μM, P2X3)) were applied to DRG neurons in a random order through a gravity-driven 12 barrel perfusion system for 10s following 30s ECS wash between stimuli to evoke AP generation under current clamp mode. Both agonist solutions were made up in pH 7.4 ECS from respective stock solution (1mM capsaicin stock in 100% ethanol, Sigma-Aldrich; and 5 mM α,β, mATP stock in 100% ethanol). The average delta spike in response to each agonist was calculated by spike numbers (normalized by subtracting spike numbers at pH 7.4) divided by agonist application time.

A standard voltage-step protocol was applied on tested neurons under voltage-clamp mode to determine the voltage-current relation^6^. Cells were held at −120 mV for 240 ms before stepping to the test potential (−50 mV to +40 mV in 10 mV increments) for 40 ms (Figure 1H). Voltage was returned to holding potential (−60 mV) for 200 ms between sweeps. Leak subtraction was applied to minimize capacitive currents. Step current density was calculated by minimum (inward) and maximum (outward) current amplitude (pA) (normalised by subtracting average baseline amplitude (100 ms) at –120 mV) dividing cell capacitance (pF). Calculated current density (pA/pF) was then plotted against corresponding step voltage (mV) as voltage-current relation and fitted in Igor Pro using a single or double Boltzmann equation.

### Ca^2+^ imaging

DRG neurons were incubated in 10 μM Ca^2+^ indicator Fluo-4 AM (diluted from a 10 mM stock solution in DMSO in ECS; Invitrogen, UK) for 30 mins at room temperature (21°C). Neurons were then washed with ECS and placed on microscope (Nikon Eclipse Tie–S, Nikon) for imaging. Fluro-4 fluorescence was excited by a 470 nm LED (Cairn Research) and images were captured by a digital camera (Zyla cSMOS, Andor) at 1 Hz with 50 ms exposure time using Micro-Manager software (v1.4; NIH).

The same ion channel agonist solutions used in electrophysiology and 50 mM KCl (to serve as a positive control) were applied on neurons following an established perfusion protocol: 10 s ECS wash following 10 s agonist application and another 90 s wash in ECS. All solutions were perfused through a gravity-driven 12-barrel perfusion system. A 3 min interval was applied to allow the neurons to return to their resting state among each perfusion.

Data analysis was carried following in house protocol^5^. Briefly, KCl positive cells and one black background were draw manually as region of interest (ROI) using ImageJ software and mean gray value of selected ROIs in sequence was extracted. Extracted data were then analyzed by lab-developed R toolbox (https://github.com/amapruns/Calcium-Imaging-Analysis-with-R) to calculate Ca^2+^ influx change (normalized to peak KCl response (ΔF/Fmax) with background subtraction) and the percentage of agonist respondent cells (cells with ΔF/Fmax value less than 0.001 and peak after 30 s were deleted manually.).

### Statistics

All figures presented were analyzed and graphed in Graphpad Prism 8 or IgorPro software unless stated otherwise. Data shown as mean ± Standard Error of Mean (SEM). Two group comparisons were carried by Student’s unpaired t-test, and percentage comparison was done by chi-square test. One-way ANOVA (Analysis of Variance) with Tukey post-hoc test was used to compare the significance among three groups. P values less than 0.05 was considered significant.

## Supplementary Tables

**Table 1:** scores of osteochondral defects in sheep by ICRS macroscopic scoring system

| Sheep # | ICRS macroscopic score (appearance of joint post mortem) |
| --- | --- |
| 1 | 3 |
| 2 | 2 |
| 3 | 2 |
| 4 | 2 |
| 5 | 3 |
| 6 | 2 |

**Table 2:** Action potential properties of sheep DRG neurons in Ctrl and OD groups. * signifies p < 0.05, unpaired t-test.

|  | Ctrl (n = 15) |  | OD (n = 16) |  |
| --- | --- | --- | --- | --- |
|  | Mean | SEM | Mean | SEM |
| Diameter ($\mu\text{m}$) | 39.5 | 2.1 | 39.5 | 1.9 |
| Capacitance (pF) | 71.0 | 10.6 | 80.5 | 11.9 |
| RMP (mV) | -48.3 | 3.5 | -46.5 | 2.4 |
| Threshold (pA) | 116.7 | 40.1 | 100.0 | 56.5 |
| Half peak Duration (HPD, ms) | 3.1* | 0.2 | 4.5 | 0.5 |
| Afterhyperpolarization duration (AHP, ms) | 73.9 | 9.6 | 82.7 | 7.0 |
| Afterhyperpolarization amplitude (AHP, mV) | 14.0 | 1.9 | 14.0 | 2.3 |
| Amplitude (mV) | 101.6 | 4.7 | 103.5 | 2.8 |

**Table 3:**
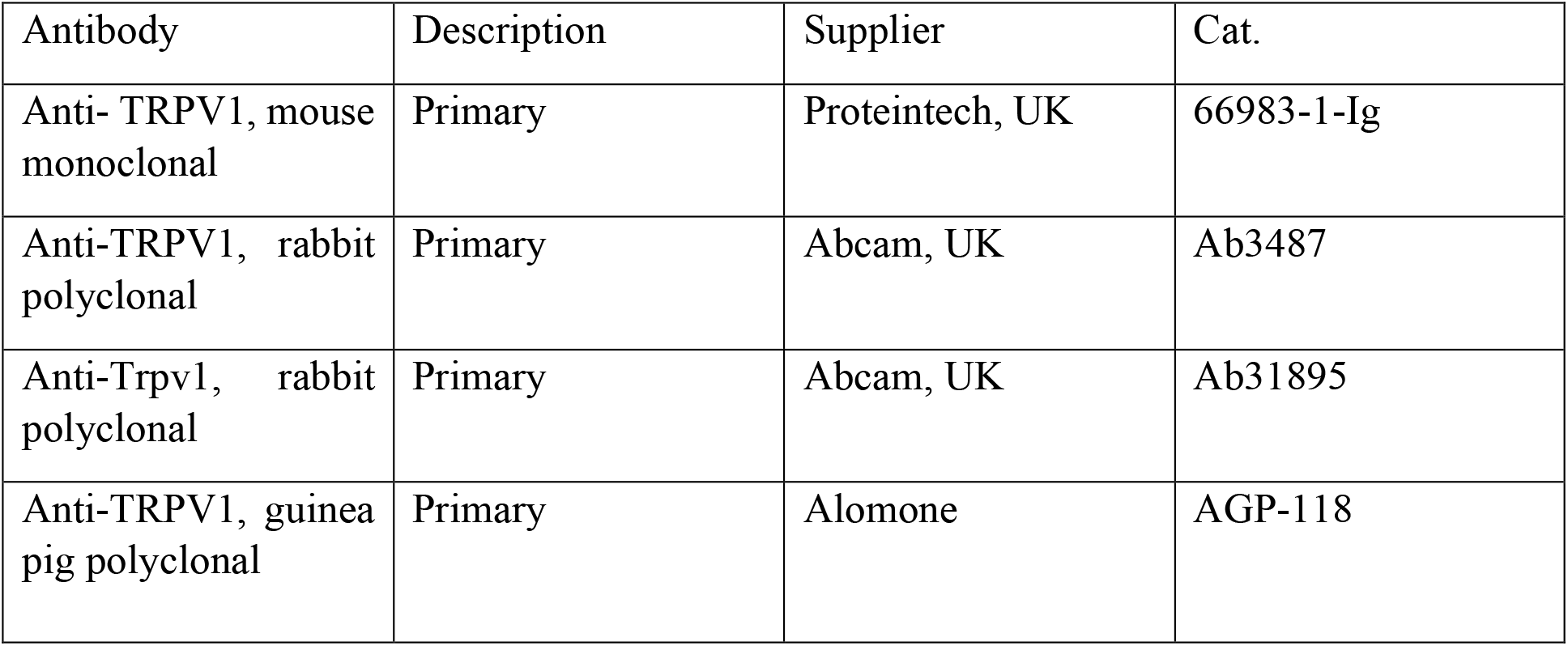
List of TRPV1 antibodies tested on sheep DRGs

| Antibody | Description | Supplier | Cat. |
| --- | --- | --- | --- |
| Anti- TRPV1, mouse monoclonal | Primary | Proteintech, UK | 66983-1-Ig |
| Anti-TRPV1, rabbit polyclonal | Primary | Abcam, UK | Ab3487 |
| Anti-Trpv1, rabbit polyclonal | Primary | Abcam, UK | Ab31895 |
| Anti-TRPV1, guinea pig polyclonal | Primary | Alomone | AGP-118 |

**Supplementary figure legend:**
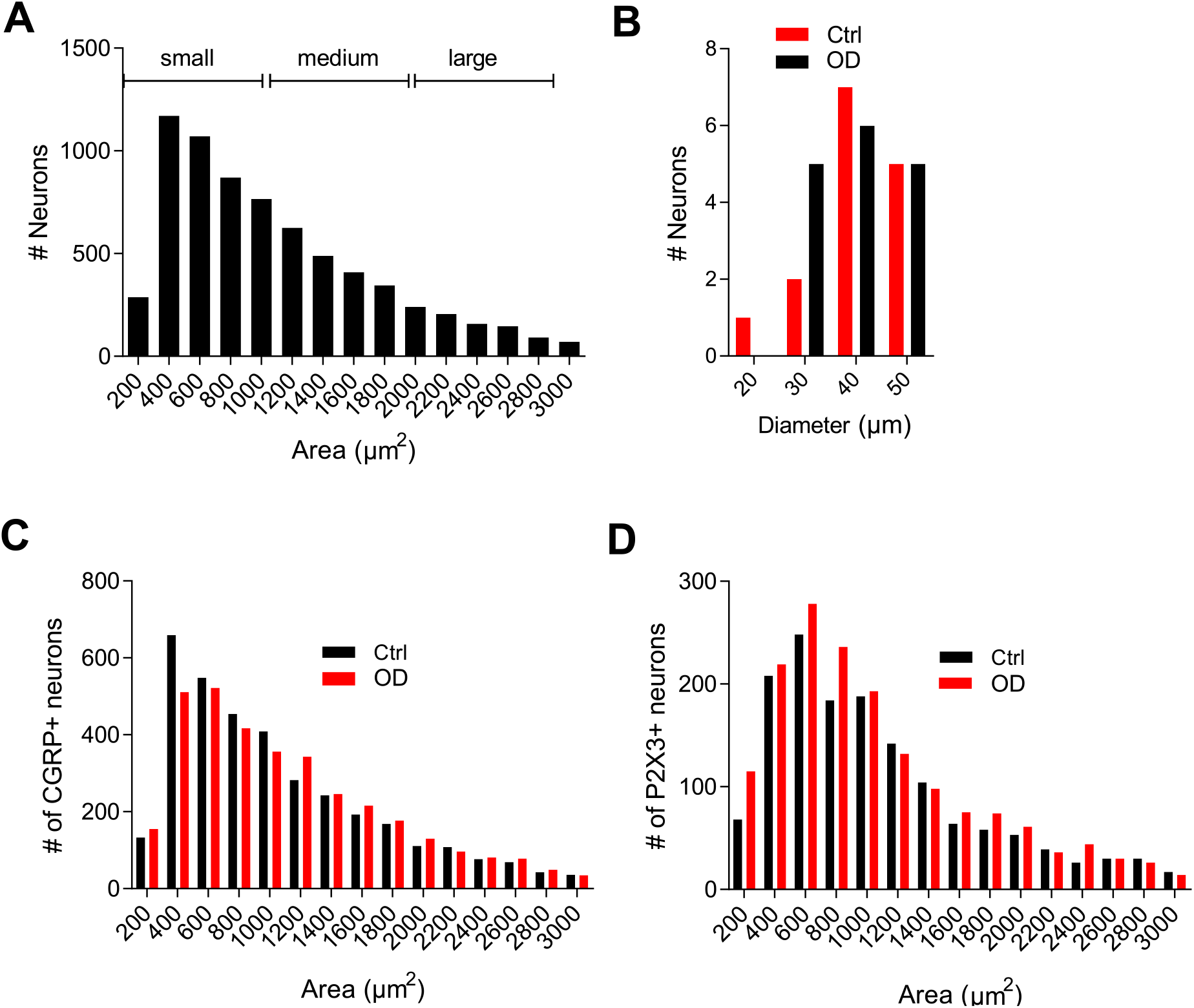
A) Histogram showing area of each sheep DRG neuron imaged from whole DRG sections and the criteria used in this article for assigning neurons into small, medium and large category. B) Histogram of neuronal diameters on which whole-cell patch clamp was performed. Histograms of cross-sectional areas of neurons stained positive by anti-CGRP (C) and anti-P2X3 (D) antibodies.

